# The Drug Repurposing Encyclopedia (DRE): a web server for systematic drug repurposing across 20 organisms

**DOI:** 10.1101/2023.03.10.532084

**Authors:** Xuexin Li, Lu Pan, Laura Sanchez-Burgos, Daniela Hühn, Oscar Fernandez-Capetillo

**Affiliations:** Science for Life Laboratory, Division of Genome Biology, Department of Medical Biochemistry and Biophysics, Karolinska Institute, 171 21 Stockholm, Sweden; Department of Medical Epidemiology and Biostatistics, Karolinska Institutet, Solna, 17165, Sweden; Genomic Instability Group, Spanish National Cancer Research Centre (CNIO), Madrid 28029, Spain; Research Institute of Molecular Pathology (IMP), Vienna BioCenter (VBC), Vienna 1030, Austria

## Abstract

The identification of new therapeutic uses for compounds via computational or experimental approaches, which is widely known as drug repurposing, has the potential to develop novel therapies with pre-existing medicines, thereby reducing the time and costs associated with drug development. Today, several data-driven methodologies have been developed leading to databases that facilitate drug repurposing initiatives. However, no approach has systematically compared drug transcriptional profiles to those from a wide spectrum of human diseases or molecular pathways. Here, we present the Drug Repurposing Encyclopedia (DRE, https://www.drugrep.org), an interactive web server covering over 198M significant drug-signature associations across 20 organisms to allow users to carry out drug-repositioning analyses. DRE consists of 12 modules covering real-time drug-repurposing for user-provided transcriptional signatures; gene set enrichment analysis (GSEA) for all available drug transcriptomics profiles; as well as similarity analyses for provided gene sets across all database signatures. Collectively, DRE provides a one-stop comprehensive solution to help scientists interested in drug-repurposing studies.

## INTRODUCTION

Drug development is a time-exhaustive and cost-intensive process with a median time to clinical development of 8.3 years over the past decade (1). These long development times may pose immediate life-threatening impacts during the emergence of novel diseases such as the pandemic outbreak of COVID-19. A fast alternative is to discover new therapeutic usages for existing medicines via computational or experimental methods, also known as drug repurposing, and which has already been proven successful in several examples (2,3). This pragmatic approach has the potential to save time and costs for drug development and for the formulation of new therapeutic hypotheses. In his regard, methodologies and databases related to drug repurposing have already emerged (2-6). One example is the Connectivity Map (CMap), which stores over 30,000 transcriptional signatures from drugs and genetic perturbations in various cell types, enabling users to run multiple analyses, such as for instance trying to find drugs triggering similar transcriptional changes and thus likely to share a common mechanism of action (4). Another potential use of CMap is to find drugs that cause a transcriptional signature opposite to that associated from a given disease, thereby being a potential treatment for this pathology (7,8). As an example, we recently used such an approach to predict drugs that could potentially counteract (or aggravate) the severity of the cytokine storm arising in severe cases of COVID-19 (9). Yet, no repository has systematically compared the transcriptional signatures associated to drugs, to those from diseases or specific signalling pathways.

To this end, we used the signatures available at the largest and most popular repository; the Molecular Signatures Database (MSigDB) (10,11), together with drug transcriptomic profiles available at the Connectivity Map (CMap). By performing a systematic comparison of these signatures, we built an interactive drug-repurposing database, named the Drug Repurposing Encyclopedia (DRE, https://www.drugrep.org), which harbours 198,648,641 drug-signature associations across 20 organisms. Besides storing these pre-computed associations, the DRE web server allows users to carry out real-time drug-repurposing analysis to compare user-provided gene signatures with those available at the DRE database, or to perform drug-gene set enrichment analyses (drug-GSEA) for provided drug transcriptomics profiles as well as similarity analyses of provided gene sets across all database signatures. In summary, DRE is the first comprehensive webserver dedicated to facilitate drug repurposing approaches based on transcriptional signatures, through in-depth assessments across molecular signatures and species.

## MATERIALS AND METHODS

### Data Collection

#### Molecular Signatures

A total of 648,825 molecular signatures were collected from the MSigDB v7.5.1 (10-12), comprising signatures from 20 organisms (an average of 32K signatures per organism) (**Table 1**). We included all 9 major molecular-signature categories from MSigDB, including: hallmark gene sets of well-defined biological states and pathways (H); chromosomal positional gene sets (C1); curated gene sets from publications, including KEGG (13) and Reactome (14) pathways (C2); regulatory target gene sets (C3); cancer-oriented computational gene sets (C4); ontology gene sets, including Gene Ontology (GO) terms (15) (C5); oncogenic gene sets (C6); immunological gene sets (C7); and cell type signature gene sets (C8) (10,11).

#### Drug Profiles

For our analyses, we used the 4,690 consensus drug profiles available at DREIMT, a drug repurposing database focused on immunomodulation (16), which are tabulated drug transcription profiles that were in turn imported from the CMap-associated Library of Network-Based Cellular Signatures (LINCS) L1000 (4). Specifically, the downloaded drug profiles were constructed from Level 3 data that derive from gene expression counts of 978 landmark genes normalized against invariant gene sets and normalized across experimental plates (4). Expression for another 11,350 genes was inferred from normalized landmark gene counts. Differential expression analysis for each drug profile was conducted and an additive linear model used to correct for bias introduced by batch and individual cell line drug responses (16). Final drug profiles are consensus transcriptional changes of the drugs across cell lines and experimental conditions (16,17).

### Web Server Construction

#### Association Analyses

Systematic enrichment analyses were performed to evaluate the similarity between drug-associated transcriptional signatures available at CMap with all the molecular signatures from 20 organisms present at MSigDB. Specifically, GSEA analyses were carried out for each of the 648,825 molecular signatures from MSigDB against 4,690 consensus drug profiles based on the adaptive multilevel splitting Monte Carlo approach (18). The preliminary estimation of p-values was based on 10,000 permutations for each analysis. Altogether, 3,042,989,250 associations were obtained across 20 organisms. Multiple testing correction was done using the Benjamini-Hochberg (BH) false discovery rate (FDR) (19), after which a final set of 198,648,641 associations with an FDR < 0.05 were considered as significant and retained to be included in the database. In any case, while all the significant 198M associations can be downloaded from the web server, to facilitate the analysis and concentrate on the most significant ones we only display those with an FDR < 0.01. The pipeline used to populate the DRE database is shown in **Fig. 1**.

**Figure 1.**
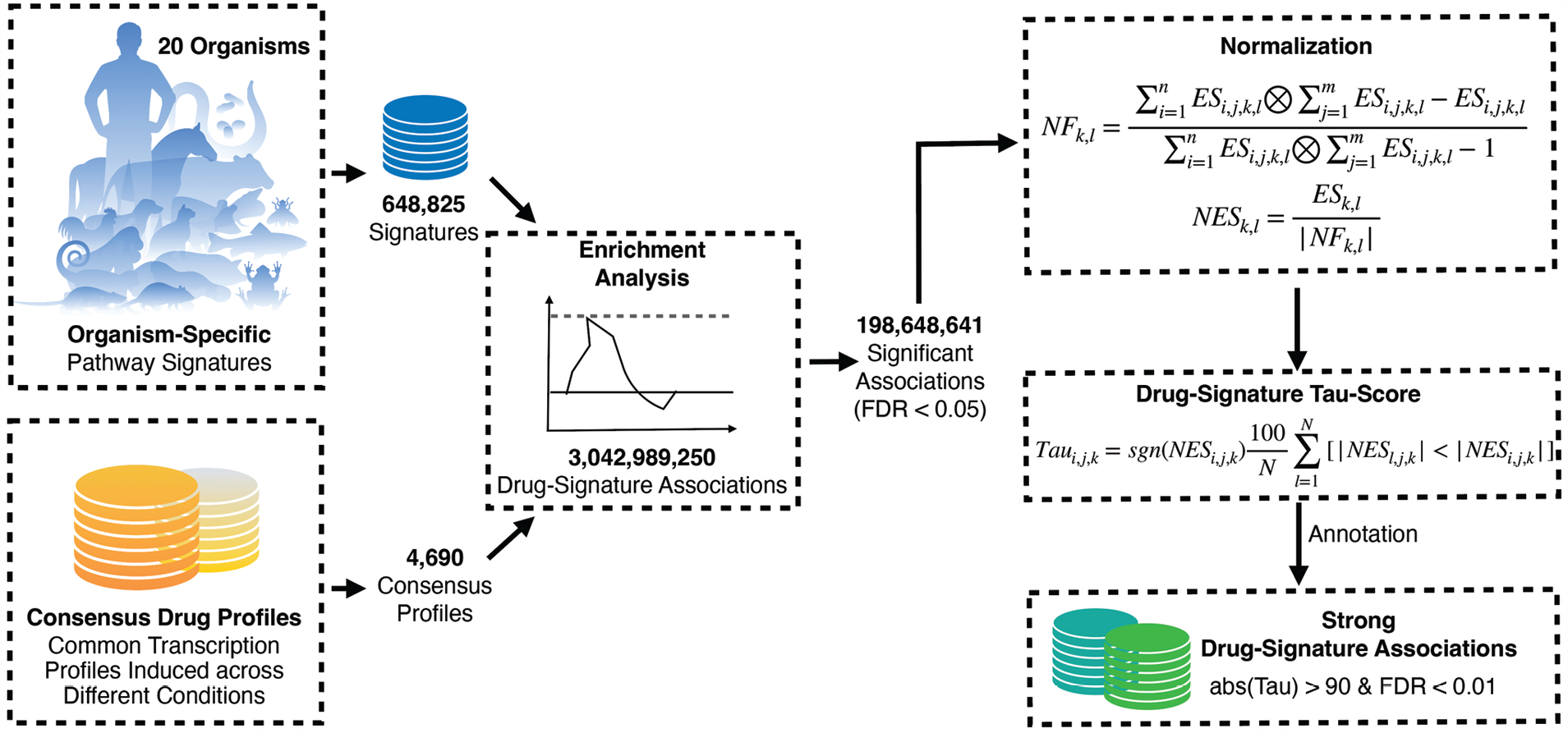
Schematic workflow of the pipeline used to select the significant drug-signature associations that build the DRE database. As mentioned, while the database displays those associations with an FDR < 0.01, an entire list of the associations with an FDR < 0.05 can also be downloaded by the user.

#### Drug Prioritization Scores

We adopted the approach by LINCS L1000 (4) and DREIMT (16) to assess the specificity of a given association of the drugs with each signature using a standardized drug prioritization Tau score (4). Enrichment scores for each molecular signature from GSEA analyses results were used to calculate Tau scores. For each organism, associations with positive or negative enrichment scores (ES) were separately normalized by the mean ES of the molecular signatures and drugs profiles by first obtaining the normalization factor NF,

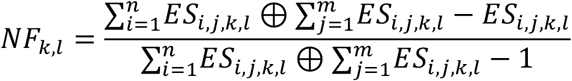

where NF represents the normalization factor for organism *k* and association *l*, being *l* = 1 for positive ES and *l* = 2 for negative ES, *i* representing molecular signatures and *j* drug profiles. The ES set for each organism *k* was divided by its normalization factor to obtain final normalized ES (NES) scores as follows,

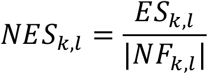

Normalized positive and negative ES values were collectively standardized for each drug profile to obtain the Tau score for each molecular signature *i* and drug profile *j*,

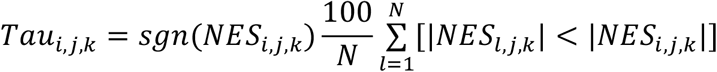

where *l* represents the *l*th NES score in drug profile *j*. The final standardized Tau scores ranges from -100 to 100. Significant signature associations were considered those with an absolute Tau score > 90 and an FDR < 0.01 (4,16).

### Customized Analysis

#### Drug Repurposing Tool

DRE includes tools to enable the comparison of user-provided gene sets against the molecular signatures from the chosen organism using GSEA (16). To optimize the process, 200 permutations for the preliminary estimation of the p-values are used for each comparison. Multiple testing corrections is done using BH-FDR and associations with an FDR < 0.1 retained for further analyses. The top 5,000 associations ranked by decreasing FDR are presented. To obtain the Tau scores for each drug-signature association, precomputed association data available at DRE are used together with the results obtained from the user-queried analysis for normalization of ES values by the mean ES of the signatures and the drugs. Normalized ES values are then standardized to derive the final Tau score for each association. The top 20 associations with the highest absolute Tau scores are finally displayed as well as available for inspection in an interactive table form.

#### Gene Set Enrichment

This tool facilitates organism-specific enrichment analysis of user-queried gene sets with the molecular signatures from MSigDB. All 648,825 signatures from a total of 20 organisms are available for these analyses. Specifically, the user should provide a two-column, tab-separated text file (with no header), containing gene symbols (column 1) and log-fold changes (column 2) arising from differential expression analyses. Based on their chosen organism, enrichment analyses are carried out to compare the queried gene set with the organism-specific molecular signatures, using 1,000 permutations for the preliminary estimation of the p-values. An FDR will be calculated and for each association, and enrichment results will be interactively displayed to the user as running scores and pre-ranked list, together with an interactive table showing the post-enrichment results.

#### Gene Set Similarity

Through this tool, users will be able to submit a specific gene signature and obtain a list of molecular signatures presenting a high concordance with the provided list (16). Upon choosing the organism, pairwise comparisons are carried out to compare the queried gene signature with all the molecular signatures from the selected organism. A Fisher’s exact test (20) is used to compute each comparison, and the Jaccard similarity coefficient (21) calculated to observe the degree of relevance for the similarity between the comparing groups (16). P-values are adjusted by BH-FDR. The top 20 most significant molecular signatures are visualized interactively, and the entire results are also available in an interactive table.

#### Drug Set Enrichment Analysis (DSEA)

We adopted the DSEA approach from (9). Briefly, instead of using gene-sets on gene-level summary statistics in a conventional GSEA procedure, drug-sets are used to conduct enrichment analysis for a set of drugs with ranked Tau scores. For these analyses, we used the fgsea method (11). Based on the mechanism of action (MOA) of each drug provided by DREIMT (16), we classified drugs into MOA categories and created a database with their associated drug signatures. Even though the aim of drug repurposing is to identify new functions for existing drugs, it is also important to know if the list of candidate drugs obtained from drug-repurposing analyses presents an enrichment for a class of drugs based on their MOA. After running the drug-repurposing tool, the DSEA tool allows the user to input the list of candidate drugs and their associated Tau scores to examine for the potential enrichment (positive or negative) of a particular class of drugs based on their MOA. Based on the GSEA method, instead of gene sets, MOA drug signatures are used and their enrichment assessed from user input data based on drug-level summary statistics. After calculating ESs, these will be presented interactively as enrichment plots for every MOA category available. An overview of the different tools available for customized analyses is provided in **Fig. 2**.

**Figure 2.**
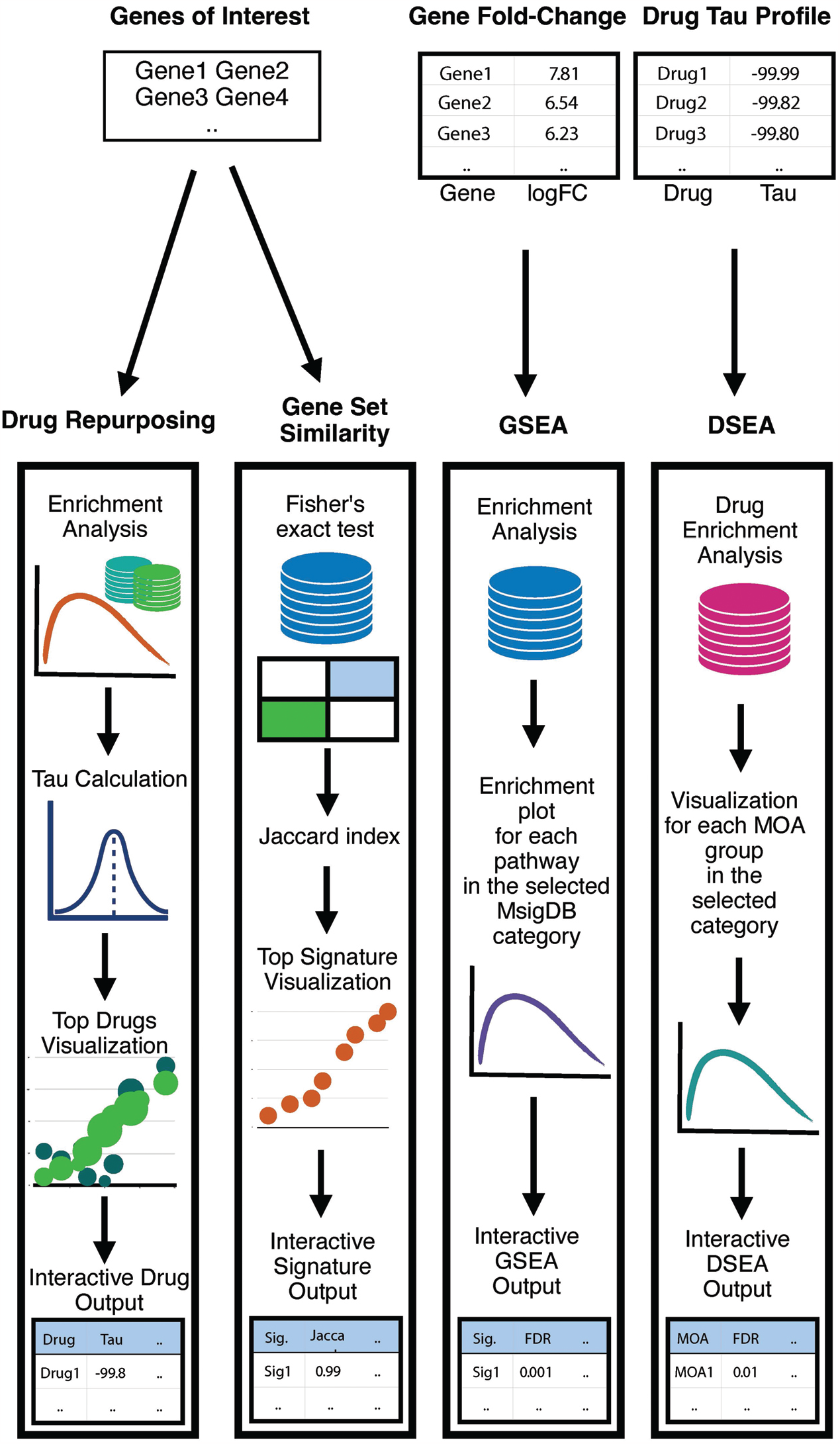
Workflow for the “Customized Analyses” modules available at DRE. For “drug repurposing” and “gene set similarity” modules, users must supply a set of genes of interest. After analyses are ran, DRE will return interactive visualization plots as well as tables with the full results. For “GSEA” and “DSEA” modules, users must supply gene-level or drug-level summary statistics, respectively, to obtain enrichment results.

## RESULTS

The purpose of DRE is a user-friendly web server that facilitates drug repurposing based on transcriptional signatures to bench scientists with a limited computational background. In this context, DRE consists of 12 modules focused on drug repurposing and the analysis of transcriptional signatures, including modules to search for similarities between drug signatures and those from specific diseases or molecular pathways; and modules for customized analyses of user provided signatures (**Fig. 3**). We here provide an overview of the different modules from the web server as well as examples to illustrate the usefulness of each of the approaches.

**Figure 3.**
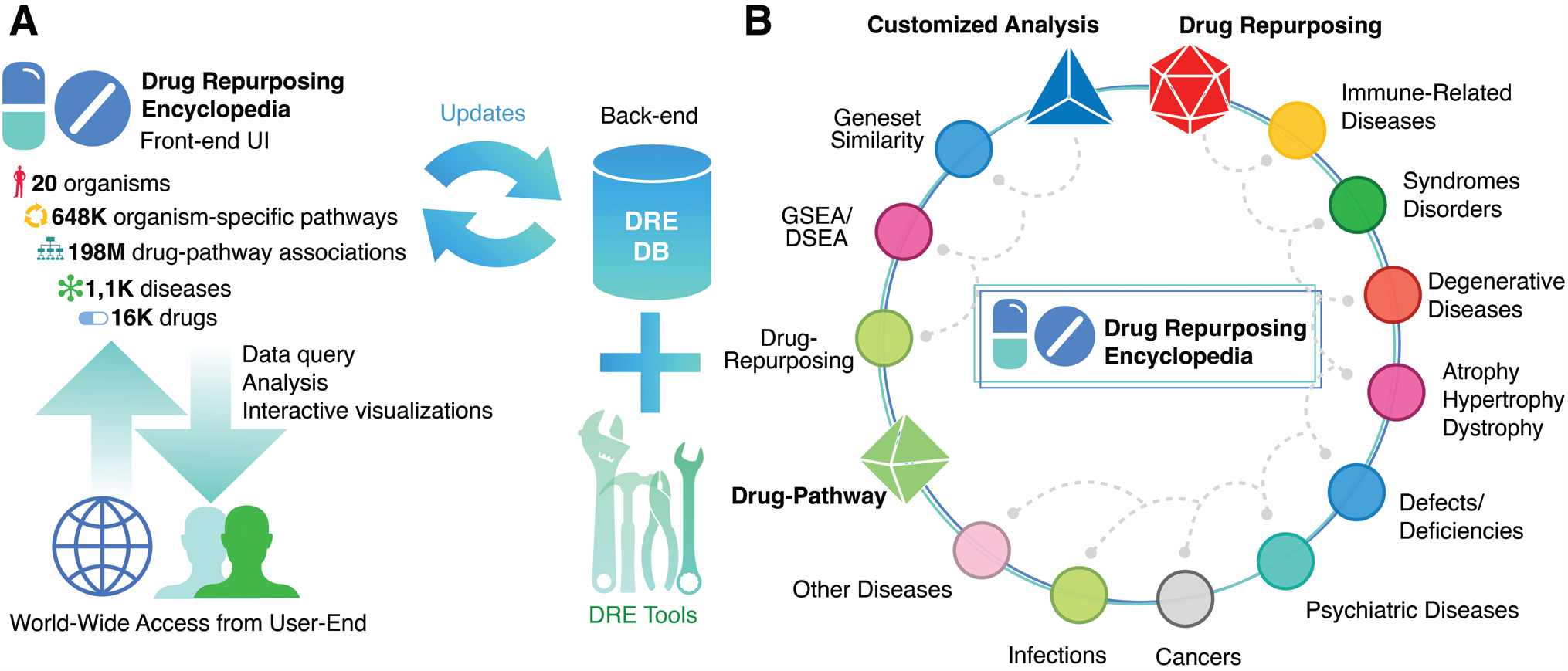
Overview of the DRE web server. (**A**) The user-interface (UI) of DRE and its backend structure, illustrating the number of organisms, signatures and drugs available for inspection. DRE can be accessed freely by users through the internet providing a large dataset of precomputed associations of transcriptional signatures for data exploration as well as several tools for analyses on user-provided datasets. The backend will be consistently updated. (**B**) A snapshot of all the modules available at DRE, including drug-repurposing associations for 9 types of diseases, associations between drugs and molecular pathways, and several tools for customized analyses on user-provided datasets.

## 1. Modules of the database for drug-disease and drug-pathway associations

### 1.1 Drug-Repurposing

Within this module, users can explore for drug-associations to 9 types of diseases ([1] immune-related diseases, [2] syndromes-disorders, [3] degenerative diseases, [4] atrophy/hypertrophy/dystrophy, [5] defects/deficiencies, [6] psychiatric diseases, [8] cancers, [8] infections and [9] other diseases). Within each module, options are available to select an organism and a specific disease. For instance, if a search is carried out after selecting “Huntington’s disease”, an autosomal-dominant, neurodegenerative disease caused by an abnormal CAG expansion (polyQ) in the first coding exon of human Huntingtin (*HTT*) (22) in the “degenerative diseases” module for Homo Sapiens, the server will display drugs with transcriptional signatures that mostly resemble or oppose the one associated to the disease. As examples, benzamil and prostaglandin-A1 have high negative Tau scores (**Fig. 4A)**, suggesting that the drugs could help to counteract the transcriptional alterations associated to HD. Consistently, benzamil has been shown to facilitate the degradation of polyQ-containing mutant HTT (23,24) and prostaglandin-A1 has a reported neuro-protective effect in preclinical models of HD (25).

**Figure 4.**
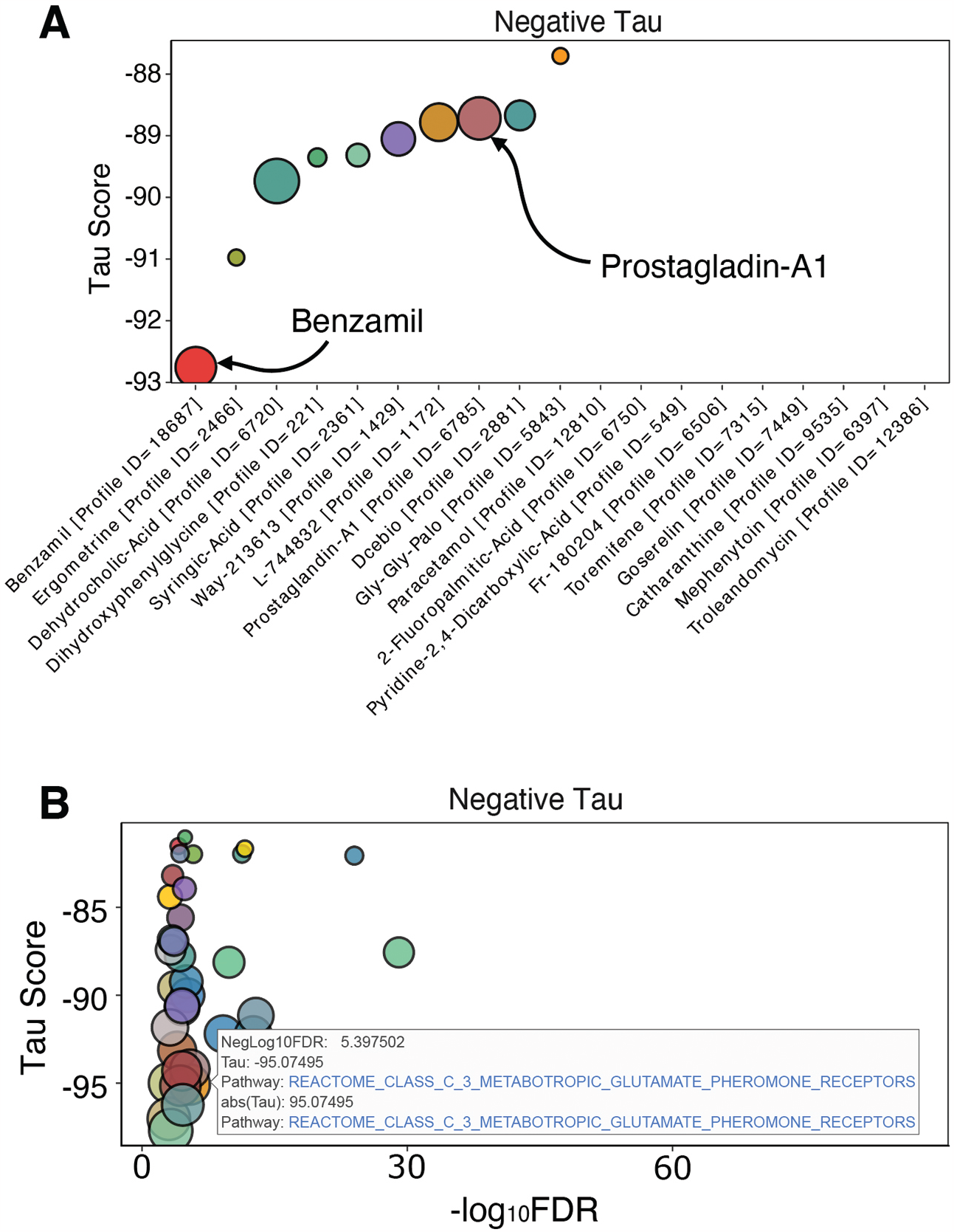
Examples of different association modules on DRE. (**A**) Example output of the “Drug Repurposing” module after searching for HD as a disease in Homo sapiens. Highlighted drugs have already been explored for the treatment of the disease (**B**) Example output of the “Drug-pathway” module after searching for the molecular signatures with signatures associated to that of latrepirdine, a glutamate receptor antagonist. As expected, there is a highly significant negative association with glutamate/pheromone receptors.

### 1.2 Drug-Pathway

The “drug-pathway” module can be used to query for a selected drug and identify molecular pathways that share similarities with its associated transcriptional signature. Once again using HD as an example, a search for the glutamate inhibitor latrepirdine, showed a negative Tau score of -95.1 with the glutamate/pheromone receptor pathway signature, that is known to play a role in several neurodegenerative conditions. In fact, latrepirdine has been used in clinical trials for HD (26,27) where it showed beneficial effects on cognition (27) (**Fig. 4B**). An interactive plot showing the top pathways with the highest positive and negative scores is shown, together will a table with the full list of pathways that are significantly associated to the drug.

## 2. Modules for customised analyses of user-provided datasets

### 2.1 Drug Repurposing

The drug repurposing tool allows users to identify drugs that could be related to their pathway of interest based on user-provided gene signatures. To this end, users need to supply a specific gene set from their experiments, as a list of space-separated gene symbols. An interactive plot displaying the drugs with the most significant association to the provided gene set is shown to the users, together with an interactive table displaying the complete results of the analysis. As an example, using as input the gene signature associated to HD from the KEGG pathway database (13), we obtain several drugs that have been already explored for the treatment of HD in cell and animal models such as gliclazide (28) and Ebselen (29) (**Fig. 5A, B**).

**Figure 5.**
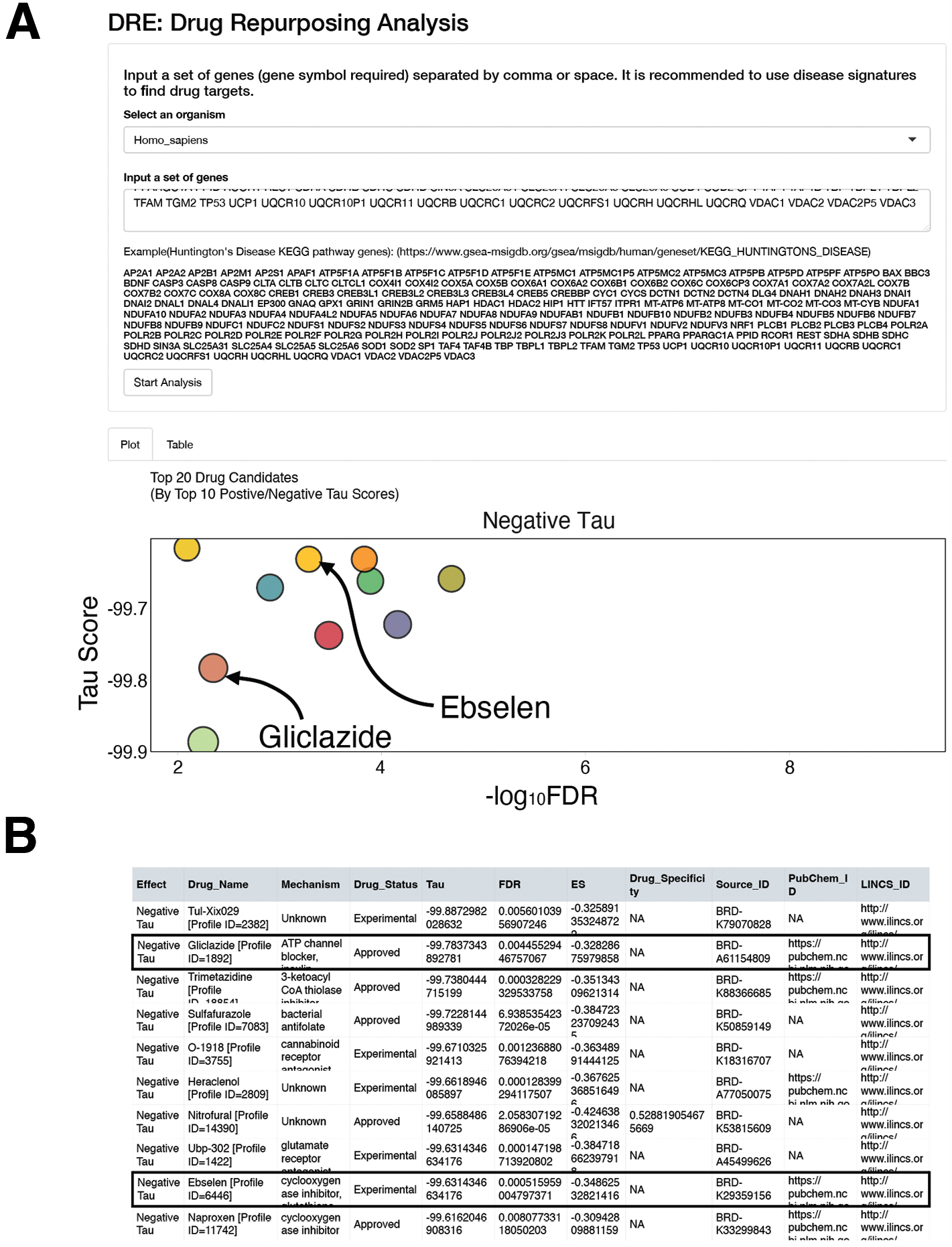
Output from the “Drug-Repurposing” analysis. (**A**) Interactive output from the “drug repurposing” analysis using the signature associated to HD available at KEGG in homo sapiens. Selected examples of drugs with a significant negative association and known to be related to HD therapy are highlighted. (**B**) Snapshot of the full results table from the previous analysis that are provided, once again highlighting the same two drugs selected in (**A**).

### 2.2 GSEA

For GSEA, users need to supply a two-column tab-separated file of differentially expressed genes (column 1 containing gene symbols, and column 2 the associated fold-change). Supplied data are then compared against a specific signature category selected from MSigDB, and an interactive GSEA plot is shown to the user, together with an interactive table containing the full results. To demonstrate the usefulness of this module, we used our own transcriptional data from U2OS cells expressing a fragment of the exon 1 from HTT harbouring an expanded track of 94 glutamines. After selecting the MSigDB C2 category to conduct the GSEA analysis, the results show a significant association of our signature to pathways related to mitochondrial biogenesis, which is already known to be affected in HD patients and consistent with our own experimental findings using this cell line (**Fig. 6A**) (30).

**Figure 6.**
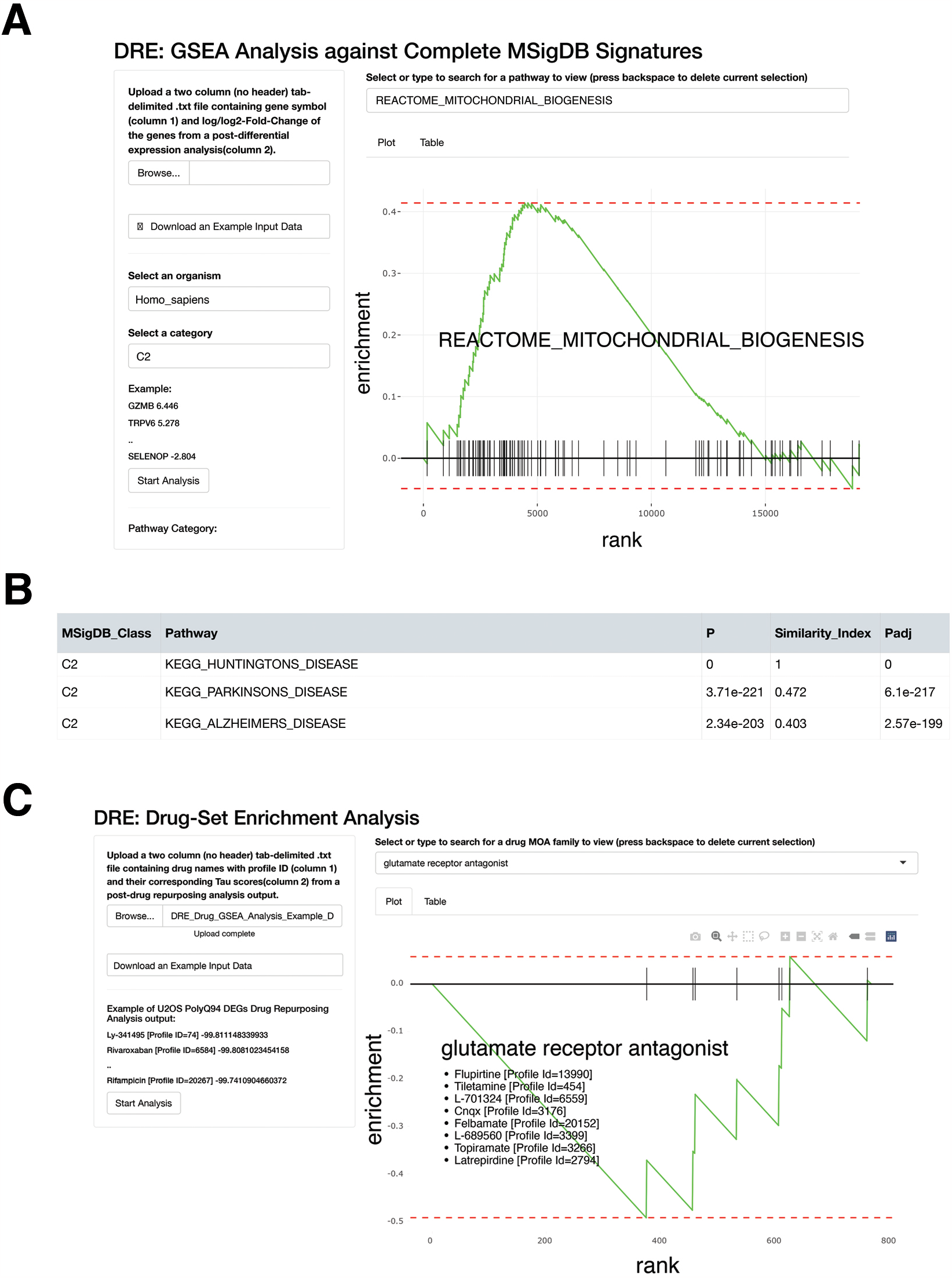
Example outputs from customized analysis at DRE. (**A**) A GSEA plot from the “Reactiome_Mitochondrial_Biogenesis” pathway after running the analysis with a signature arising from a cellular model of HD. The analysis is run against all MSigDB signature set and interactive GSEA plots can be visualized for the selected significantly associated pathways **(B)** Results from a “gene-set similarity” analysis using the HD signature from KEGG as input. As expected, the highest association is with the input signature, followed by signatures from other neurodegenerative diseases (**C**) Output from a DSEA analysis, using an input the results from the drug-repurposing analysis from the cellular model of HD mentioned in **Fig. 5**. Interactive DSEA plots can be visualized for the selected significantly associated drug classes, such as the “glutamate receptor antagonist” shown.

### 2.3 Gene Set Similarity

The gene set similarity analysis measures the degree of similarities between two gene sets. In this case, the supplied set is compared against the entire molecular signatures available at MSigDB for a selected organism. Pathways with statistically significant similarity are displayed interactively, ranked by significance. For instance, selecting the hallmark gene set associated to HD from KEGG, the analysis reveals that the highest correlation is with the same disease signature from MSigDB, along with signatures from other neurodegenerative disease pathways (**Fig. 6B**).

### 2.4 DSEA

The purpose of running DSEA is to identify potential drug classes, rather than specific drugs, that could be associated to a given signature. To use this tool, the user has to supply a two-column tab-separated table with drug names (column 1) and their corresponding Tau scores (column 2) arising from a previously ran drug-repurposing analysis. The MOA database containing a list of drug signatures grouped on the basis of their MOA, is then used to conduct an enrichment analysis with the user-supplied data set. Using the results from the drug repurposing analysis made from the transcriptional signature of U2OS expressing the polyQ-containing exon 1 of HTT, the analysis returns an enrichment for glutamate receptor antagonists among the drugs with negative Tau scores (**Fig. 6C**). As mentioned above, glutamate signalling is well-established to have a detrimental effect on HD progression, supporting the use of the antagonists for the treatment of the disease (31,32).

## DISCUSSION

We here present DRE, a one-stop solution for drug-repurposing based on transcriptional data that was built by systematic combinatory exploration of two of the largest and most popular molecular signatures repositories: CMap, for drug-associated transcriptional profiles, and the signatures available at MSigDB covering 20 different organisms and hundredths of pathways, diseases or signalling routes. The effort led to a collection of more than 198M significant associations which are readily available for inspection at DRE, thereby providing an easy first-inspection tool for those interested in drug repurposing initiatives. Besides highlighting specific drugs, DRE is also useful as a hypothesis generator for bench scientists without a computational background that are interested in performing drug-repurposing and identifying potential pathways that are related to their disease of interest. The user interface of DRE has been designed to be easy for users to manipulate and also optimized for high-volume data traffic. Altogether, we are confident that DRE can become a useful repository to help efforts from the scientific research community related to drug repurposing.

## ACKNOWLEDGEMENTS

The authors want to thank Dr. Fatima Al-Shahrour for valuable input on this work.

## AVAILABILITY

The DRE webserver and its database are freely available at https://www.drugrep.org. Codes for this study have been deposited at https://github.com/xuexinliki/DRE.

## TABLE AND FIGURES LEGENDS

Table 1. Summary statistics of pre- and post-processing of the molecular signatures of organisms from the MSigDB.

